# Elucidating the role of multiple feedback loops in regulating stem cell decisions

**DOI:** 10.1101/2024.09.25.615049

**Authors:** Razeen Shaikh, Gregory T. Reeves

**Affiliations:** Department of Chemical Engineering, Texas A&M University, College Station, Texas 77843, United States; Interdisciplinary Program in Genetics and Genomics, Texas A&M University, College Station, Texas 77843, United States

**Author notes:** To whom correspondence should be addressed: Gregory T. Reeves, 100 Spence Street, Texas A&M University, Chemical Engineering Dept College Station, Texas 77843 MS 3122.

## Abstract

Stem cell decisions are regulated by a complex network of gene regulatory pathways that determine the reproductive health of the tissue. The *Drosophila* ovarian germline is a well-characterized model system which facilitates visualizing stem cell behavior in its native environment to attain a systems-level understanding of the stem cell dynamics. The asymmetric division of the Germline Stem Cells (GSCs) forms two daughter cells–a self-renewed GSC and a differentiated Cystoblast (CB). The highly conserved Bone Morphogenetic Protein (BMP) pathway ensures growth and maintenance of the GSCs, but is downregulated in the CBs, to allow for differentiation. BMP signal transduction upregulates *dad* and represses Fused, both of which are negative regulators of the BMP pathway. Moreover, these regulatory mechanisms operate on a system of two cells which remain connected during a portion of the cell cycle. We developed a biologically-informed mathematical model of multi-compartment GSC division to investigate the dynamic roles Dad and Fused play in determining cell fate. We found that Dad optimally controls the BMP signal transduction to enable GSC homeostasis and differentiation. In *dad*^KO^ mutants, GSCs were more likely to divide symmetrically. Our work identifies the synergistic role of Dad and Fused rendering robustness to stem cell division.

## 2 Introduction

Stem cells determine the reproductive health of a tissue and can help diagnose and treat diseases, including infertility and ovarian cancer. The exact mechanisms through which stem cells influence tissue growth and development is unknown since their decisions are regulated by complex and intertwined signaling networks. The germline stem cells (GSCs) in the *Drosophila* ovary remain functional throughout adulthood making them a great model system to perform a systems-level investigation of stem cell decisions in their native environment. Moreover, the *Drosophila* germline has fewer redundancies in the signaling pathways which regulate stem cell decisions compared to their vertebrate homologs. Despite these advantages, there are only a few mechanistic models of the signaling pathways to help discern the underlying rules of stem cell decisions (Harris, et al., 2011; Pargett, et al., 2014; Sardi, et al., 2021; Tian, et al., 2012; Xia, et al., 2012).

The germline stem cells reside at the apical tip of the germarium (Figure 1A) and undergo asymmetric division to form two daughter cells: a self-renewed GSC and a differentiated cystoblast (CB). The terminal filament, cap cells, and anterior escort cells form a stem cell microenvironment, or “niche,” enabling GSC growth and maintenance. The GSC daughter cell that moves away from the niche assumes a differentiated fate. However, the GSC cell division cycle has a duration of ∼12-15 h during which the GSC and the presumptive-CB (preCB) remain connected to each other for a duration of ∼1.5 h (Figure 1B).

**Figure 1:**
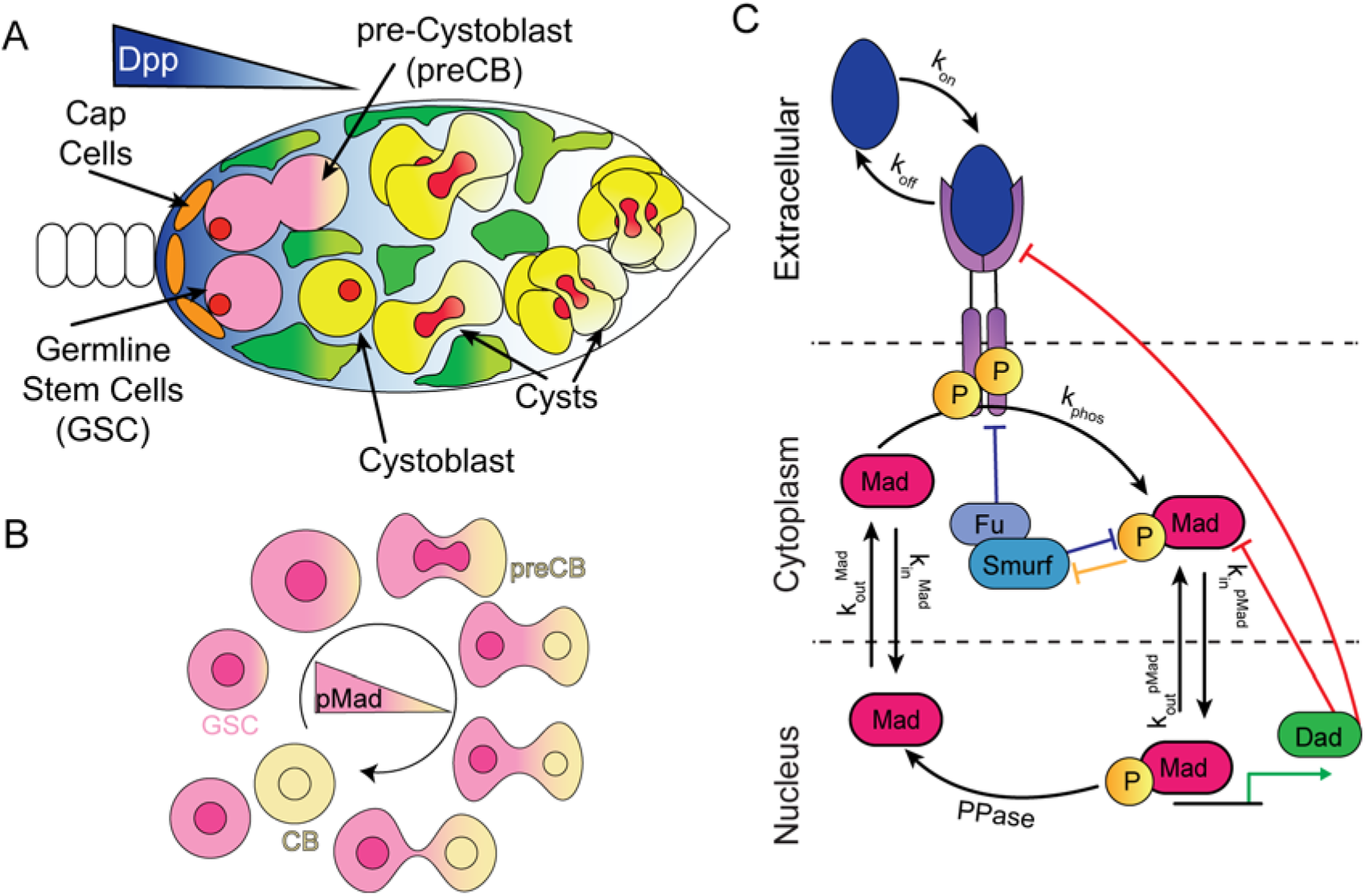
BMP signaling in the Germline Stem Cell niche. **(A)** Illustration of the *Drosophila* Germarium**. (B)** The GSC division cycle **(C)** The BMP signaling network in the GSCs and preCB/CB.

Several signaling pathways, including the JAK/STAT, Wnt, and BMP pathways, synergistically regulate the growth, maintenance, and differentiation of GSCs (Chen and McKearin, 2005; Xie, 2013; Zhang and Cai, 2020). However, the highly conserved BMP pathway plays a dominant role in influencing stem cell decisions (Fisun Hamaratoglu and Markus Affolter and George, 2014; Harris and Ashe, 2011). Decapentaplegic (Dpp), the primary *Drosophila* BMP ligand (homolog of vertebrate BMP2/4), is produced in the cap cells and diffuses to the GSCs and preCB/CBs; consequently, the two cells experience different levels of the extracellular ligand (Ridwan, et al., 2022; Sardi, et al., 2021).

Dpp forms a heterodimeric complex by with the secondary ligand Glass Bottom Boat (Gbb), which binds to the transmembrane receptors Thickveins/Punt (Tkv/Punt) or Thickveins/Saxophone (Tkv/Sax), forming a heterodimer-heterotetrametric complex. This lig- and-receptor complex triggers the intracellular SMAD signal transduction pathway by phosphorylating Mothers against Dpp (Mad; *Drosophila* homolog of vertebrate Smad1/5/9) (Harris and Ashe, 2011; Wilcockson and Ashe, 2019). Phospho-Mad (pMad) binds to transcription factor Medea (Med; *Drosophila* homolog of vertebrate Smad4) and facilitates its translocation to the nucleus where it can bind to DNA and regulate the expression of numerous genes, including the inhibitory-SMAD homolog Daughters against Dpp (Dad; *Drosophila* homolog of vertebrate Smad6/7) (Casanueva and Ferguson, 2004; Wilcockson and Ashe, 2019). Dad downregulates BMP signaling, forming a negative feedback loop (NFL), by either degrading pMad or deactivating the receptors (Li, et al., 2017). The BMP signal transduction represses the Fused/SMAD specific E3 ubiquitin protein ligase (Fu/Smurf) complex, which is itself known to downregulate BMP signaling, forming a positive feedback loop (PFL) (Xia, et al., 2010; Xia, et al., 2012). The downregulation of the BMP pathway in the CB is essential to allow for the expression of the differentiation factor, *bag of marbles* (*bam)*, which remains repressed in the GSCs.

To ensure precise and robust maintenance and differentiation of the GSCs, the BMP pathway must be differentially activated such that the pMad/Med complex is available to the GSCs but depleted in the CBs. However, the GSC and preCB share an interface, which enables active transport of proteins between the two cells during a 1.5 h time window in the ∼12 h of the cell division cycle. A quantitative understanding of the BMP signaling network, which is currently lacking, would help discern the design principles of stem cell regulation over a short range. To overcome this limitation and drive experimental discovery, we developed a dynamic mathematical model of the BMP network in the GSC as it undergoes cell division. We constrained the variable model parameters by evaluating the ability of the model predictions to infer qualitative biological experiments in the literature. We verified that our model conforms to the system behavior reported in the literature. This allowed us to investigate possible dynamic roles of both the negative feedback element through Dad and the positive feedback element through Fused in determining cell fate.

Consistent with previous literature (Xia, et al., 2012), we found that Fused (in the PFL) can make the system bistable, such that in the *wildtype* system CB-like levels of Dpp are low enough to allow Fused expression, and consequently, differentiation. On the other hand, in *dad*-knockouts (*dad^KO^*), the same Dpp levels result in higher pMad/Med concentration, which, in some cases, maintains repression of Fused, even in the CB. This suggests that, in *dad*^KO^ mutants, GSCs might be more likely to divide symmetrically. Our simulations also indicate that Dad and Fused make the GSCs and CBs robust to perturbations in Dpp levels, respectively. Further, we inferred that Dad determines the rise time of Fused, which could enable temporal control of GSC differentiation. Together, our model identifies the synergistic role of the negative feedback element through Dad and positive feedback element through Fused in rendering robustness to the stem cell maintenance and differentiation.

## 3 Methods

### 3.1 Model architecture

To simulate the role of BMP signaling and negative feedback through Dad in the GSC, we first formulated a two-compartment GSC model with nucleus and cytoplasm (Figure 2A; see Section 3.1.1 for description of model components and interactions and the Supplement for model equations). To mathematically formulate the process of GSC division, resulting in the formation of two asymmetric daughter cells, we extended our equations to a four-compartment GSC cell-division model of the BMP network in which the GSC and the preCB experience differential levels of Dpp, and the interface connecting the two cells shrinks as they divide (Figure 2). We used the steady state of the two-compartment GSC model as initial conditions to the GSC cell-division model (Figure 2A; see Section 3.1.1 for description of model components and interactions and the Supplement for model equations). However, the two-compartment GSC model differs from the four-compartment GSC cell-division model in total volume and in Dpp signaling across two cells. To account for these differences, we simulated the BMP network in the four-compartment GSC cell-division model to calculate the pseudo-steady state concentrations of BMP pathway components without altering the mass transfer rate of proteins between the two cells. Finally, we use the pseudo-steady state concentrations as initial conditions to simulate the GSC cell-division model with a dynamic mass transfer rate to model interface shrinkage (Figure 2B).

**Figure 2:**
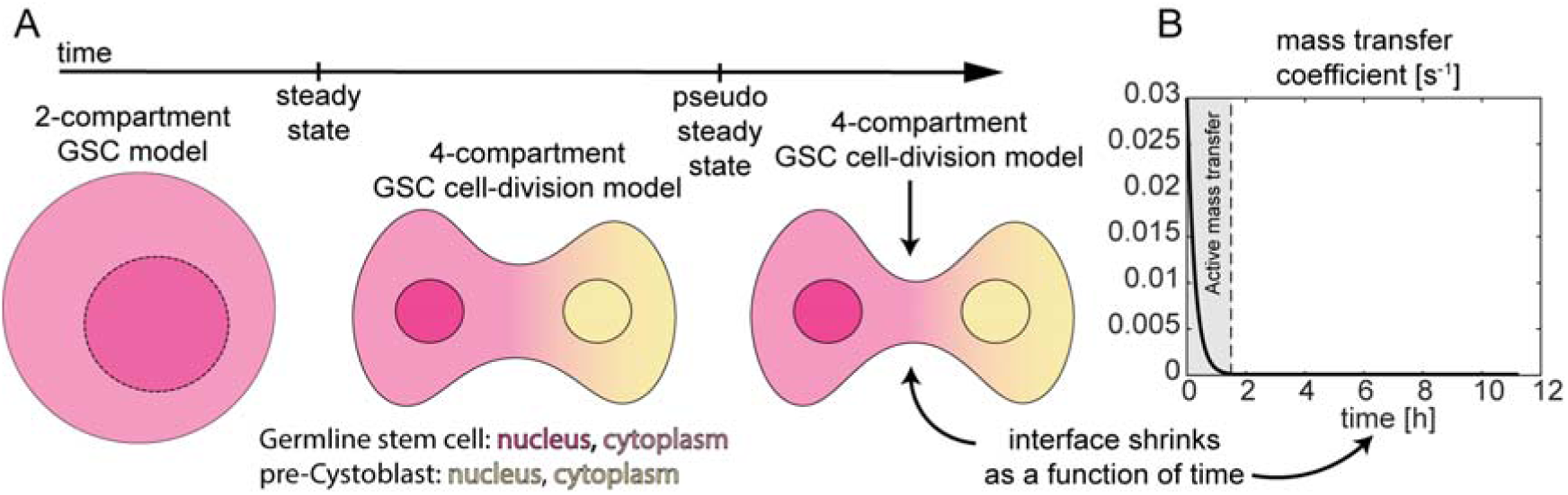
Model Architecture. **(A)** A GSC model and the four compartment model with a time-invariant mass transfer rate precedes the GSC cell-division model. **(B)** The mass transfer coefficient of BMP pathway components in the GSC cell-division is an exponential function of time.

#### 3.1.1 The BMP/Mad signal transduction network

An external stimulus of Dpp activates the signal transduction pathway by phosphorylating the transcriptionally inactive Mad to the active pMad. To reduce the model complexity, the heteromeric ligand complexes of Dpp/Gbb or Dpp/Gbb (Harris and Ashe, 2011), which activate the pathway, are formulated as a single component (Dpp), and transcriptionally active trimeric (pMad)_2_/Med complex is formulated as pMad. We allow both the inactive Mad and the active pMad to continuously exchange between the cytoplasm and the nucleus. We assume that pMad in the nucleus is transcriptionally active and upregulates Dad. We assume that Dad and Fu/Smurf are only present in the cytoplasm. Since there is no consensus in literature as to how Dad and Fu/Smurf mechanistically interact and downregulate Dpp signaling, in our model we allow them to degrade both the active pMad and active Tkv receptors(Harris and Ashe, 2011; Liang, et al., 2003; Murakami, et al., 2003; Podos, et al., 2001; Xia, et al., 2010; Xia, et al., 2012).

It is known that pMad inhibits Fu, but the mechanistic interaction between them is unclear in literature. To allow for ambiguity, we investigate two modes of inhibition, one in which pMad translationally represses Fu and in the second, pMad enhances the rate of degradation of Fu. The two models will be referred to as:

1. Repression Model: pMad translationally represses Fu.
2. Degradation Model: pMad enhances degradation of Fu.

The repression model has a dominant number of biologically-informed traits, which is why for most of the results in this paper we report the performance of the ‘Repression Model’. Refer to the Supplementary Section 4 for more details on the performance of the ‘Degradation Model’.

#### 3.1.2 The GSC model

The GSC model has two compartments: the cytoplasm and the nucleus.

A unit Dpp input activates the signal transduction pathway.

We use this model to understand how Dpp signaling, and Dad negative feedback regulates GSC maintenance, and we use the steady state from this model as an initial condition to the GSC cell-division model.

#### 3.1.3 The GSC cell-division model

The GSC cell-division model is a system of two connected cells and their nuclei. The model is simulated for a duration of 12 h, during which there is active transfer of entities for about 1.5 h through the interface. We assume that each compartment is well mixed throughout the simulation duration. The core network architecture of the Dpp signaling pathway remains the same in the GSC model and in GSC cell-division model. As above, the GSC experiences a unit concentration of Dpp, while the concentration of Dpp in the preCB is either set to 0.1 or determined by calculating the concentration of Dpp at which the bifurcation diagram of active pMad is bistable.

At the beginning of the simulation, the two-connected cells are assumed to be at a pseudo steady state. To determine the initial condition of this model, the GSC cell-division model is run to steady state with the initial conditions of the GSC model, with a constant mass transfer coeffi-cient. Then the mass transfer coefficient is reduced dynamically which is modeled as an exponentially decaying function.

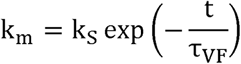

The coefficients k_s_ and *τ*_s_ are based on FRAP and long-time-course imaging by Sardi et al., 2021 and Villa-Fombuena et al., 2021, respectively (Sardi, et al., 2021; Villa-Fombuena, et al., 2021). We simulate the model for a duration of 12 h.

### 3.2 Deploying the model architecture

We simulate two genotypes with our models, the *wild-type* (*wt*) and *dad*-knockouts (*dad^KO^*). To simulate the *wild-type* model, we first determine the steady state of the GSC model and then the concentration of Dpp in the preCB. The concentration of Dpp in the preCB is assumed to be the mean of the two limit points for parameter sets that exhibit bistability, otherwise, it is set to 0.1 units. (See supplementary note 2.3 and 3.4 for a sensitivity analysis on the choice of Dpp_CB_)

To simulate *dad^KO^*, we set the scale factor value in the Hill equation for *dad* expression to be a large number (10^4^), essentially knocking it out (see Supplement for model equations).

To model GSC growth and division from a single cell, we assume that the two connected cells are at pseudo-steady state. The steady state of the GSC model was used as an initial condition for a GSC cell-division model with a time-invariant mass transfer rate. The steady state from this simulation was then used as an initial condition for the GSC cell-division model with a time-varying mass transfer rate.

### 3.3 Computing response features

#### 1. The bifurcation diagram

To compute the bifurcation curve with Dpp as the bifurcation variable, we implemented pseudo-arclength continuation on the GSC model (Supplementary Section 5). When bistability was present, we took the mean of the Dpp values at the two limit points to determine the concentration of Dpp in the preCB.

#### 2. Sensitivity coefficient (Φ)

The sensitivity coefficient is the fractional change in the response variable with respect to the fractional change in the input variable when the system input is subjected to a small perturbation (*δ*). In this study, we determined the change in steady state concentration of pMad in the GSC model and the GSC cell-division model, with respect to the change in the input Dpp level. It is mathematically formulated as:

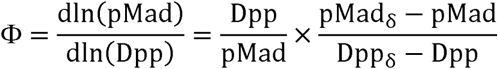

Here, pMad_δ_ is the steady state concentration when the Dpp input is increased by δ to Dpp_δ_. A unit Dpp concentration was perturbed by δ = 10^-3^ units to calculate the sensitivity coefficient.

#### 3. Rise time (t_rise_)

The rise time is defined as the duration of a response to reach 95% of its steady state value (Shaikh, et al., 2024). We calculated the rise time using a MATLAB function stepinfo from the control systems toolbox, on the dynamic profile of pMad or Fu/Smurf.

### 3.4 Parameter screens

#### 3.4.1 Unbiased screen

We generated 5 × 10^4^ parameter sets in an unbiased screen by sampling 20 or 21 scaled parameters in ‘logspace’ with bounds in the range (10^-2^,10^2^). This screen resulted in several parameter sets with unrealistic behavior. For example, some parameter sets had a high pMad dephosphorylation rate paired with a low Tkv activation rate, which resulted in permanent saturation of Fu production, which, in turn, forced pMad to be nearly zero, even in the presence of high Dpp levels. Therefore, we categorized parameter sets as “non-biologically-informed” and “biologically-informed” based on the criteria listed in the next section on “Categorizing Parameter Sets” (see also Figure 2). The “biologically-informed” parameter sets (1371 and 121 out of 5 × 10^4^ parameter sets for the Repression and Degradation models, respectively) were then used to perform a directed screen.

#### 3.4.2 Directed screen

The initial, unbiased screen had 5 × 10^4^ parameter sets, which is insufficient for a model with 20 or 21 parameters with a wide range on bounds. The unbiased screen resulted in a number of parameter sets with biologically unrealistic behavior. Therefore, to screen through biologically-informed behavior, we performed a “directed screen” by using the biologically-informed parameter sets from the unbiased screen to constrain the parameter space. We implemented strategies often used in evolutionary algorithms (“recombining” and “mutating” the best sets) to generate new parameter sets likely to result in “biologically informed” behavior (see Supplementary Section 4). The recombination strategy involves generating new parameter sets by recombining existing sets according to Supplementary Equation 33. The mutation strategy involves applying a non-isotropic Gaussian mutation using a separately defined strategy parameter to perturb parameter values and generate new parameter sets (see Supplementary Table 7). To implement these strategies, we randomly selected about half the parameter sets and performed recombination and mutation on them and repeated this process until we generated 5 × 10^4^ and 5 × 10^3^ parameter sets for the Repression and Degradation Model, respectively (Figure 2). See supplementary Section 2 for more details.

### 3.5 Categorizing Biologically-informed parameter sets

Quantitative experimental data to constrain model predictions is highly limited. The only data available is duration of the cell division cycle (Villa-Fombuena, et al., 2021) and the mass transfer rate between the GSC and the preCB (Sardi, et al., 2021), which we already incorporated in our model. To constrain model behavior, despite the lack of any *a priori* estimates of the biophysical parameters, we imposed several loose constraints based qualitative observations reported in the literature. These constraints were formulated to discard only a minimal number of parameter sets to avoid overfitting and drawing incorrect conclusions from the model.

The criteria are listed below:

#### 1. Trivial parameter sets

To check the validity of a given parameter set, we evaluate for the fold change in the concentration of pMad, Fu/Smurf, and Dad concentration between the GSC and CB at steady state in the GSC cell-division model. Parameter sets which result in a fold change of less than than 1.05 (arbitrarily selected to identify parameter sets for which model allows for a minimal dynamic change) are discarded as trivial.

#### 2. The Concentration of Dpp in the preCB

Based on the image intensity levels reported in Wilcockson and Ashe, 2019 (Wilcockson and Ashe, 2019), we estimated that the difference in concentration of Dpp between the GSC and CB might be approximately 10-fold. In our normalized model, the concentration of Dpp in the GSC is fixed to unity, so the concentration of Dpp in the CB was expected to be approximately 0.1 units. In our simulations, if bistability was detected, concentration of Dpp in the CB was chosen to be the average of the Dpp levels at which limit points were found in the *wildtype* bifurcation curve. Parameter sets for which this number was at least 0.05 units were classified as biologically-informed.

#### 3. The concentration of pMad in the CBs should be greater than Dad

Based on *lacZ* staining (Casanueva and Ferguson, 2003) we expected the concentration of Dad to be low in the CB. Model predictions which allow Dad in the CB to be higher than pMad in the CB do not align with the observed experimental data and were not classified as biologically-informed.

#### 4. The concentration of Dad should fall to the background level in the CB

Similarly, we expect the normalized level of Dad in the CB to be low (Casanueva and Ferguson, 2003). In our normalized GSC and GSC division model we implement this constraint by retaining only those parameter sets for which the normalized level of Dad in the CB is less than 0.05 units.

#### 5. The Concentration of Dad in GSC should be higher than Fu/Smurf

Since Dpp signaling in the GSCs allows for Dad expression and Fu/Smurf repression, parameter sets which predict a higher concentration of Fu/Smurf than of Dad in the GSC were not classified as biologically-informed.

#### 6. The concentration of Fu/Smurf in the CB should be higher than GSC

Based on the literature (Xia, et al., 2010; Xia, et al., 2012), we imposed a constraint such that, to be classified as “biologically-informed,” parameter sets must generate a Fused concentration in the CB which is at least greater than 1.5 times its concentration in the GSC.

## 4 Results

### 4.1 A model of BMP/Mad signaling in GSCs and CBs

In the GSC, the extracellular BMP ligand, Dpp, initiates the Smad signaling pathway. In our mathematical model, we treat extracellular Dpp levels as an input, and we model the activation of the transmembrane receptor Tkv followed by the phosphorylation of Mad to pMad and the nuclear import of both species. Nuclear pMad is assumed to be transcriptionally active, enabling *dad* expression and activating the negative feedback loop. Cytoplasmic pMad downregulates the Fu/Smurf complex, and, when present, the Fu/Smurf complex ubiquitinates pMad, forming a positive feedback loop. The mechanism through which pMad downregulates Fu/Smurf is unclear, so we evaluated two scenarios: (1) translational repression of Fu/Smurf (2) pMad mediated enhanced degradation of Fu/Smurf. We call these models “Fu/Smurf Repression Model” and “Fu/Smurf Degradation Model”, respectively.

During cell division, the GSC polarizes such that one of the GSC daughter cells, the presumptive-Cystoblast (preCB), receives a lower concentration of Dpp than the other GSC daughter cell. We model this system as two cells, GSC and preCB, which share a dynamically shrinking cytoplasmic connection. Throughout this time, the input Dpp level for the GSC is 1, while the Dpp input for the preCB is equal to 0.1 (for parameter sets that lack bistability) or is determined by the limit points on the bifurcation curve (for bistable parameter sets). The entire BMP signaling pathway is simulated in the GSC as well as the preCB, and the proteins Mad, pMad, Dad and Fu/Smurf can dynamically move between the two cells subject to mass-transfer considerations. Additionally, within each cell Mad and pMad can translocate into and out of the nucleus.

As the parameter values in these models are unknown, we initially performed an unbiased parameter screen with wide bounds in the (10^-2^, 10^2^) to determine the possible model behaviors. We imposed biologically-informed constraints on these sets (see Methods) and found that the majority (84% for Repression and 92% for the Degradation Model) of screened parameter sets resulted in non-biologically informed behavior, such as pMad levels in the GSC being lower than in the CBs (see Methods) and were discarded. After screening through all the constraints, we found that 2% and 0.04% of the screened parameter sets in the Repression Model and Degradation Model, respectively, met the criteria to be “biologically-informed” (Figure 3B).

**Figure 3:**
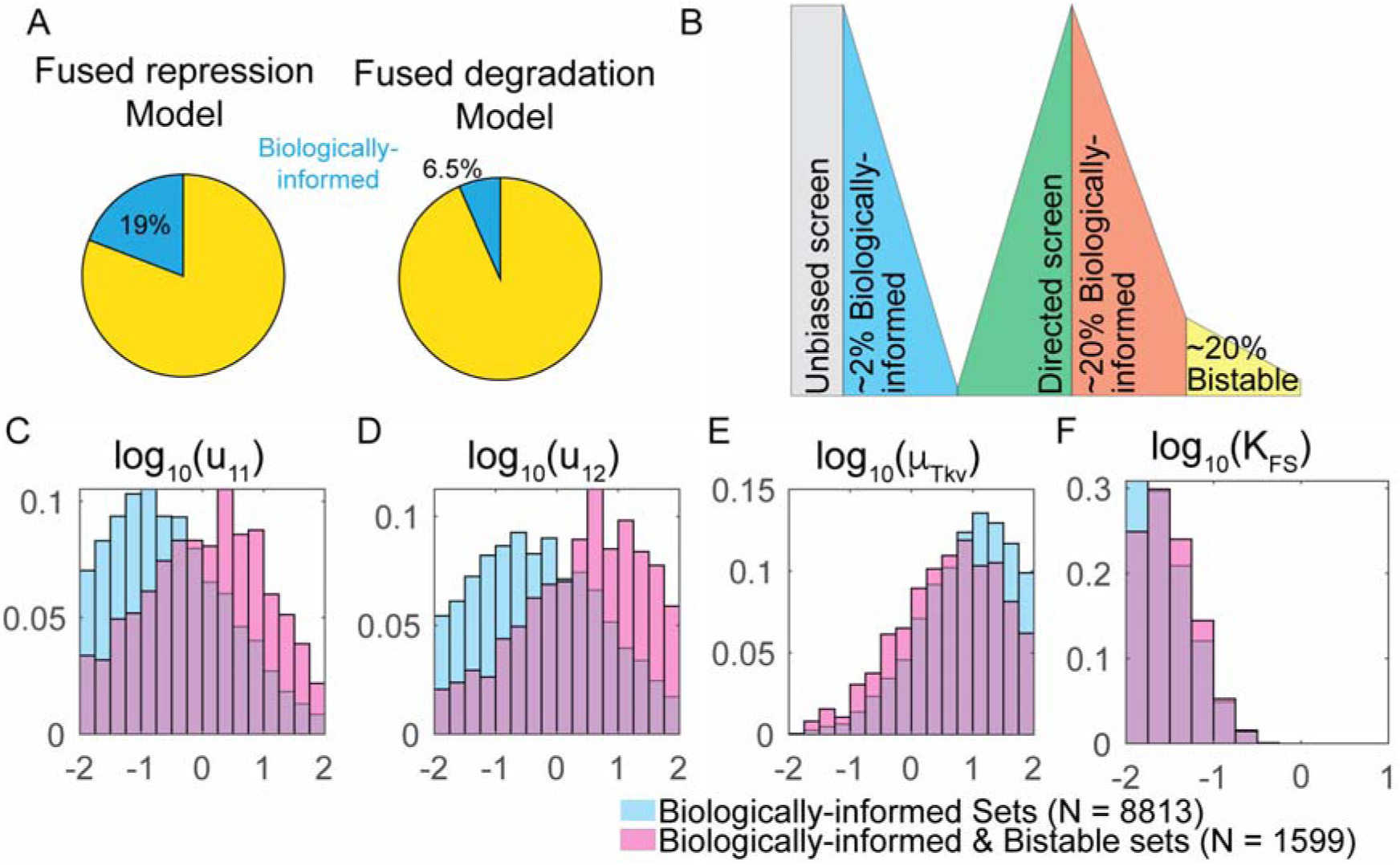
Parameter trends for biologically-informed and bistable parameter sets for the Fused repression model. **(A)** Relative abundance of biologically-informed parameter sets in the Fused repression and degradation model. **(B)** A flow diagram to illustrate the distribution of biologically-informed sets for the Fused repression model in unbiased and directed screens. **(C-F)** Selected parameter distributions predicted by the Fused repression model.

Using these biologically-informed sets, we performed a directed screen to further explore the model landscape and screen possible model behaviors. The directed screen of the Repression Model resulted in 18.8% biologically-informed parameters, whereas only 6.5% for the Degradation Model (Figure 3A). Based on this, we focused on the Fu/Smurf Repression Model (see the Supplement for more information on the Fu/Smurf Degradation Model).

The Fu/Smurf Repression Model has 20 variable parameters including synthesis rates, kinetic rate constants, Hill coefficients and scaling constants. It is difficult to measure most of these parameters *in vivo*, while some are entirely inaccessible. However, with our biologically-informed model, we were able to visualize the parameter ranges over which the model allows for biologically informed behavior for individual parameters. Of the 20 parameters (21 for Degradation Model), the rate at which Fu/Smurf degrades pMad and Tkv, the ratio of the synthesis to degradation of Tkv and the Hill coefficient of Fu/Smurf repression have the strongest effect on determining the validity of the parameter set. The distribution of these parameters is as shown in Figure 3C-F, for biologically-informed and bistable sets. Please refer to Supplementary Section 3 and 4 for the distribution of all the other parameters for both the Repression and Degradation Model.

### 4.2 Negative feedback through Dad ensures the robustness of BMP signaling in the GSC

The purpose of Dpp signaling in GSCs is to repress the differentiation factor *bam*, ensuring GSC self-renewal. As such, it seems plausible that Dpp signaling need only be higher than the threshold required to repress *bam*, making it unclear why a negative feedback inhibitor is required. Therefore, we analyzed the role of Dad in GSCs. We hypothesized that the Dad negative feedback loop is responsible for rendering the system robust, which is a common function of the negative feedback (Jermusyk and Reeves, 2016). To determine the role of Dad in ensuring robust maintenance and differentiation, we used our GSC model to calculate the robustness of the response variable (nuclear pMad) with respect to perturbations in the level of the input (Dpp), assayed by the sensitivity coefficient (see Methods). For all the biologically realistic parameter sets in the parameter screen, we compared the sensitivity coefficient in the *wild-type* GSC to that of the *dad^KO^*. We found that, on average, the GSC becomes more sensitive when *dad* is knocked out (Figure 4A). This is true not just on average, but also for each individual parameter set: knocking out *dad* decreased the robustness in greater than 89% of the biologically-informed parameter sets (Figure 4B).

**Figure 4:**
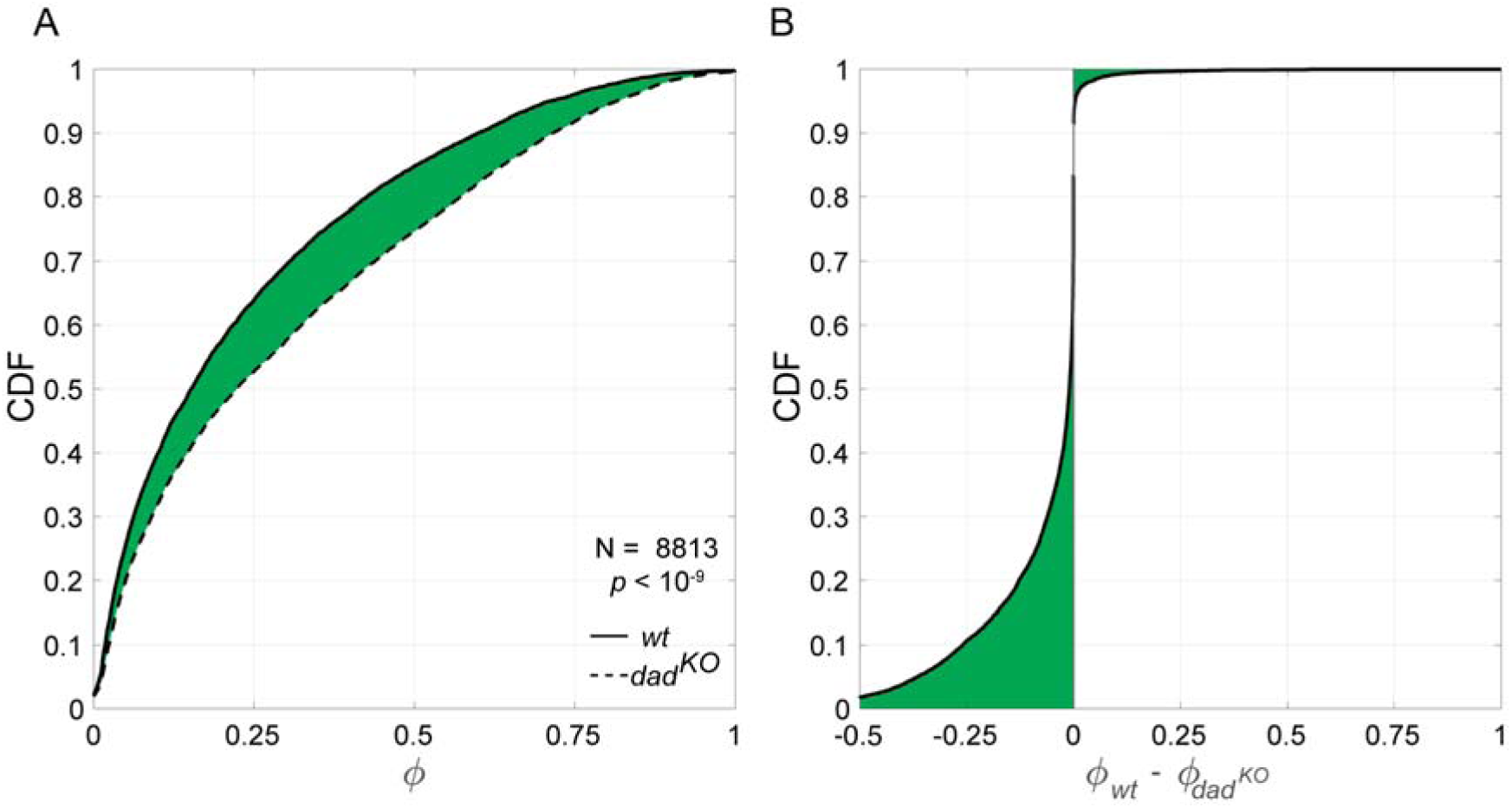
Dad renders robustness to the GSCs. The sensitivity coefficient Φ of all parameter sets **(A)** and change in sensitivity coefficient for individual parameter sets **(B)** in *wild type* and *dad^KO^* backgrounds.

### 4.3 Dad determines Fu/Smurf rise time in the CBs

Besides increasing the robustness of output with respect to input, another common role for NFLs is to speed up the system response. In the context of the GSC, the rise time may affect critical biological variables, such as the division period. Therefore, we simulated the GSC cell-division model (Figure 2A) for a duration of 12 h. Initially, mass transfer between the two cells was set to be maximal, such that they were essentially well-mixed with each other. Over the course of the first 1.5 h, the mass transfer rate decayed exponentially to zero (Figure 2B). The simulated dynamics during the 1.5 h active mass transfer period indicated that, during this stage, the concentration of pMad dips only slightly in the GSC, while it falls significantly in the preCB (Figure 5A), establishing a clear pMad difference between the two cells, as observed previously (Sardi, et al., 2021). Using the GSC cell-division model, we analyzed rise time, a performance metric for response time in both wildtype and *dad*^KO^ GSCs. We found that the rise time of Fu/Smurf in the preCB is higher in *dad^KO^* simulations than in *wild-type* (Figure 5B). For precise control of differentiation within the division timeframe, it is expected that the rise time of Fused should be tightly regulated. In *dad^KO^*, the concentration profile of takes significantly longer to respond and reach a steady state (Figure 5B), which is in agreement with delayed differentiation observed in *dad* mutants (Xie and Spradling, 1998).

**Figure 5:**
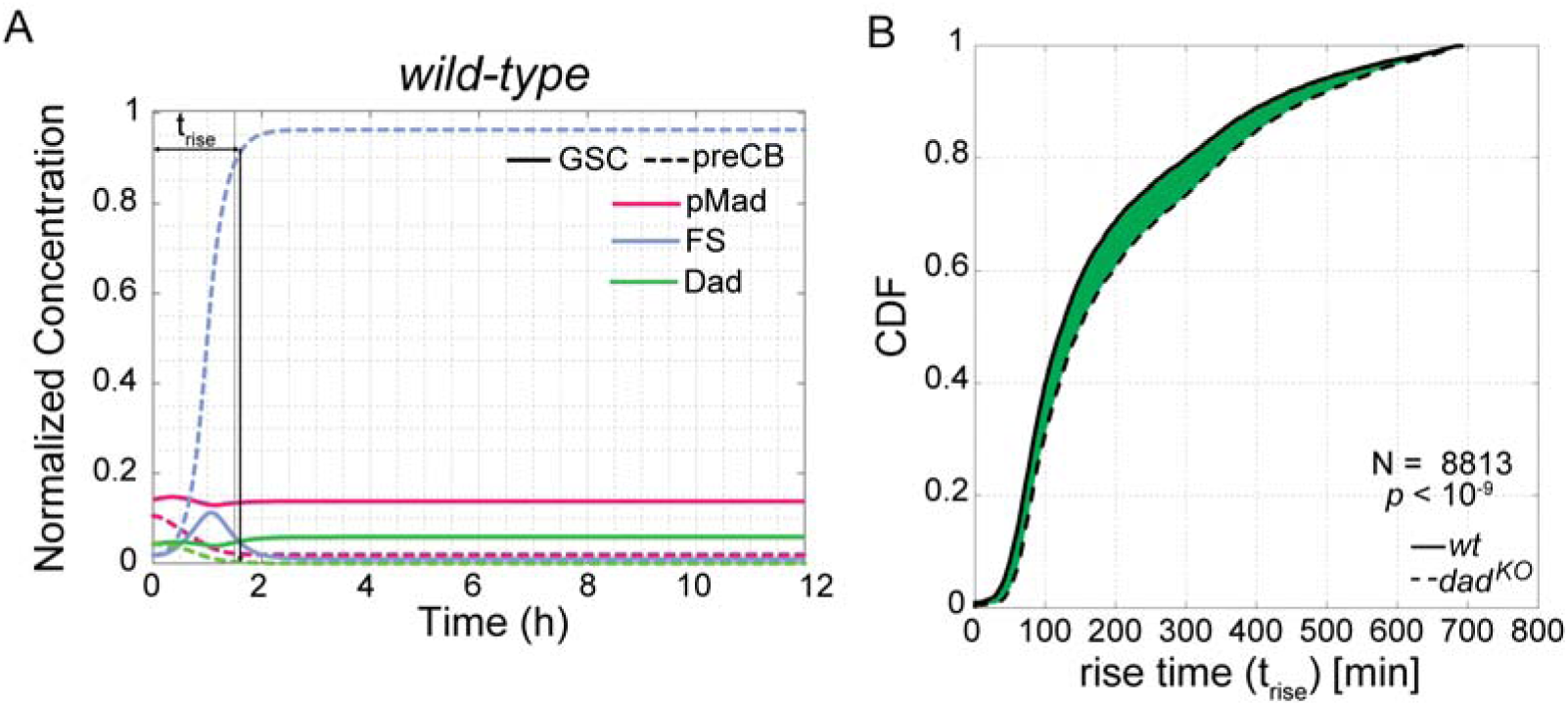
Dad regulates the rise time of Fu/Smurf. **(A)** The dynamic concentration profile of pMad, Dad and Fu/Smurf in the GSC and preCB. **(B)** The rise time of Fu/Smurf in *wild-type* and *dad^KO^* backgrounds.

### 4.4 Differentiation of the CB may be regulated by a bistable switch

In addition to negative feedback through Dad, a positive feedback loop exists in the GSC system in which Smad signaling negatively regulates Fused (Fu), a partner of the ubiquitin ligase Smurf; together, the Fu/Smurf complex causes the degradation of active Tkv receptors (Casanueva and Ferguson, 2004; Li, et al., 2017; Liang, et al., 2003; Murakami, et al., 2003; Podos, et al., 2001; Xia, et al., 2010; Xia, et al., 2012). It has previously been shown that this positive feedback loop makes the system bistable (CajaMurcia, 2006; Harris, et al., 2011; Xia, et al., 2012). Therefore, we further tested the biologically realistic parameter sets for bistability. To this end, we used pseudo-arc-length continuation on the GSC model to create bifurcation diagrams with extracellular Dpp as the bifurcation parameter (Figure 6A-C). We found that, of the biologically realistic parameter sets, 18% exhibited turning points for Dpp concentrations between zero and one, resulting in the bistability predicted for the pMad-Fu/Smurf positive feedback loop. In most of these bistable parameter sets, the bistable switch occurred at low levels of Dpp, such as those that may be experienced by the CB, suggesting that the CB can achieve either a “high” or “low” activated Tkv and pMad state (Figure 6A-C).

**Figure 6:**
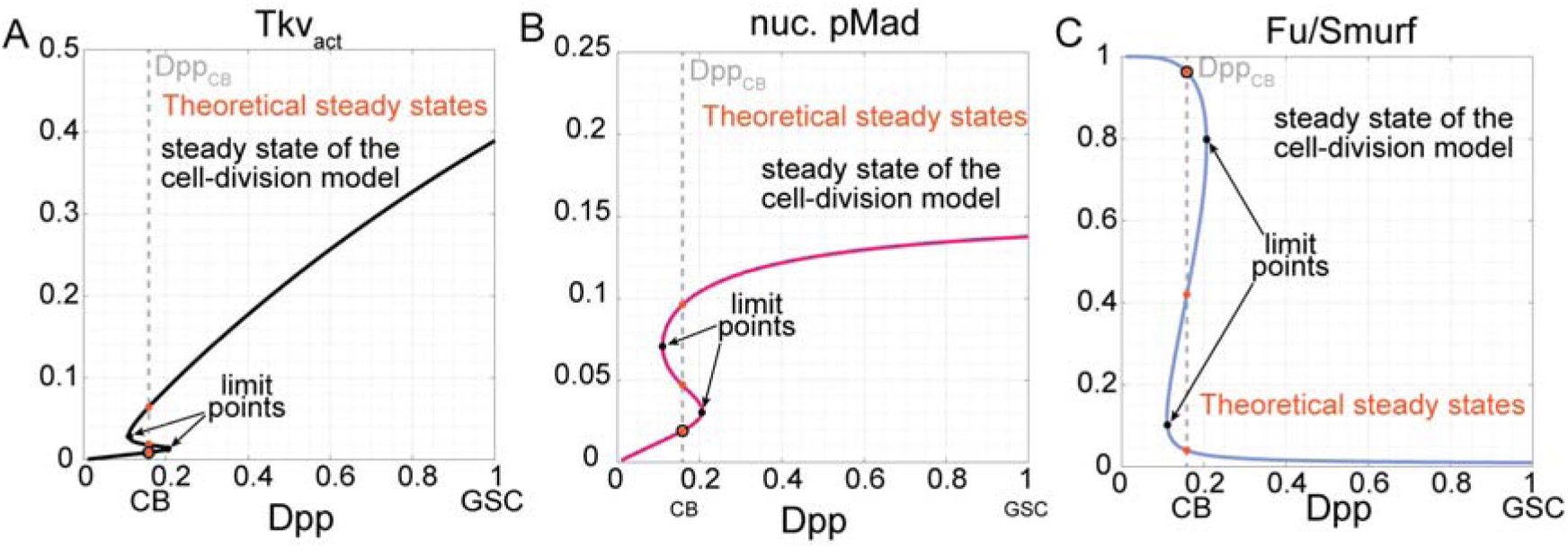
Dynamics of BMP signaling the Drosophila GSCs and CBs. **(A-C)** The bifurcation diagram for activated Thickveins (Tkv) **(A)** nuclear pMad **(B)** and Fu/Smurf **(C)**.

Given the bistability experienced by the CB, it is natural to ask whether the lower branch of Tkv_act_ and pMad concentration represents CB differentiation. Therefore, in the GSC cell-division model, we set the unknown Dpp concentration experienced by the preCB to be the average between the Dpp concentrations at the two limit points (Figure 6A-C). Under these conditions, the final steady state of the GSC cell-division model converges to the lower branch of the Tkv_act_ and pMad bifurcation curve (the higher branch of the Fu/Smurf bifurcation curve) in 10% of the parameter sets, despite having initial conditions much closer to the higher branch (Figure 6A-C). This suggests that bistability through the pMad-Fu/Smurf positive feedback loop has the capacity to control pMad and Fu expression, and thus, *bam* expression, across two-halves of a cell undergoing division. Therefore, our mechanistic GSC cell-division model helps formulate a hypothesis to infer design rules which allow the BMP pathway to regulate pMad levels in the GSCs and preCBs. However, the frequency with which the model converges to the lower branch is low (10% of the biologically-informed parameter sets that exhibit bistability), and as such, the model-informed predictions need to be validated experimentally. With the availability of new tools to track endogenous dynamics of BMP pathway elements and ability to acquire the long time-course data the model landscape can be further constrained (Hoppe and Ashe, 2021; Villa-Fombuena, et al., 2021).

### 4.5 Dad in the GSC could be necessary for the system to divide asymmetrically

To determine whether Dad plays a role in the decision for the CB to be on the upper or lower branch of the bistable system, we simulated the GSC cell-division model in *dad^KO^* ovaries. We chose only biologically-informed parameter sets for which the pMad concentration is on the lower branch of the bifurcation curve. As before, we used pseudo-arc-length continuation with Dpp as the bifurcation variable to construct bifurcation diagrams for *dad^KO^* (Figure 7B-D). Compared to wildtype, we found the bifurcation curve for the *dad*^KO^ mutant, including the upper limit point, generally shifts upward because the absence of Dad increases the level of pMad (Figure 7B-C). On the other hand, the lower branch is not significantly altered by the loss of Dad, as the steady states of the wildtype system on the lower branch already had low Dad levels (Figure 7B-C).

**Figure 7:**
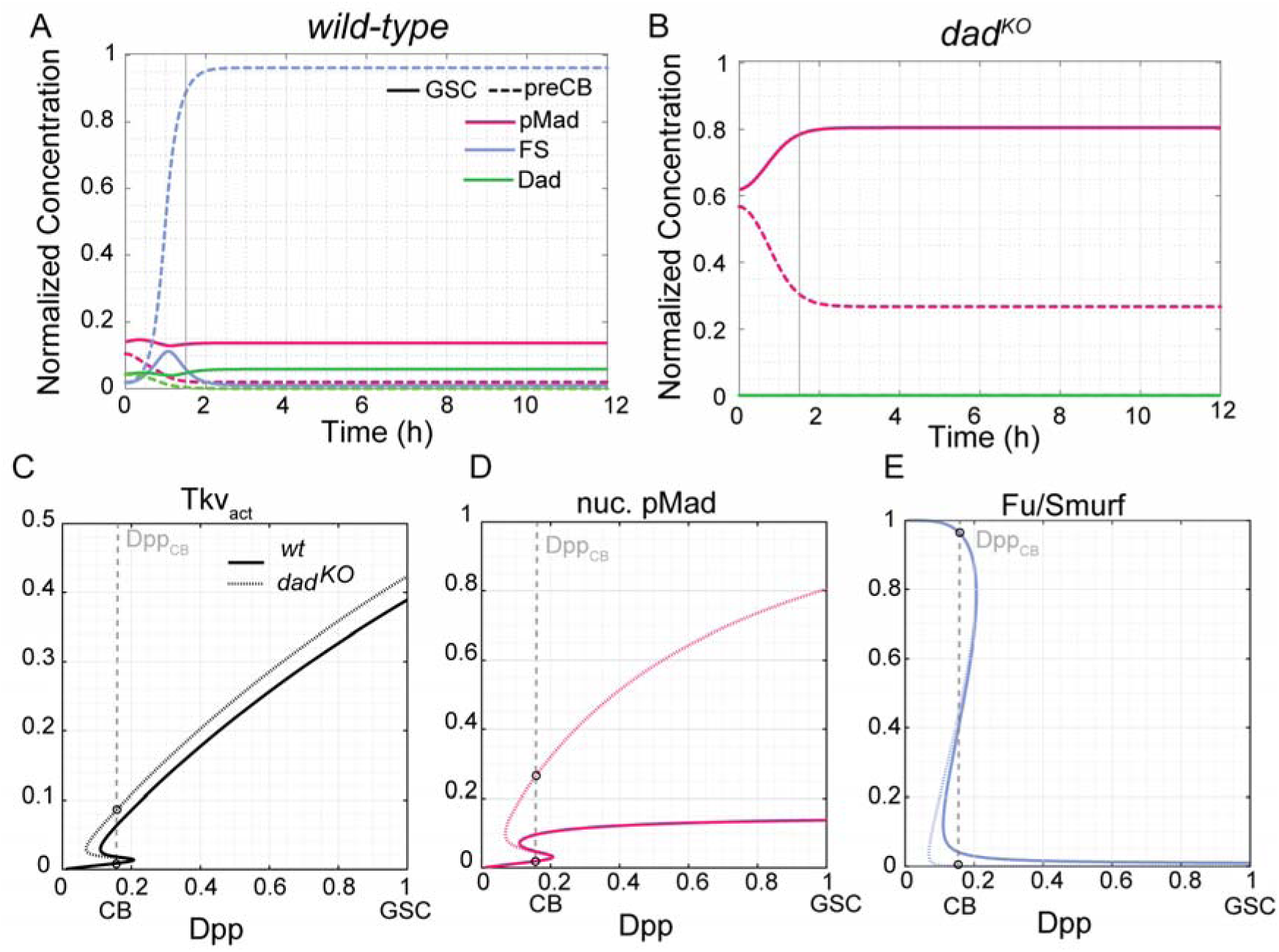
The negative feedback loop through Dad allows for asymmetric divisions of GSCs. **(A-B)** The dynamic concentration profiles of pMad, Dad and Fu/Smurf in *wild-type* and *dad^KO^* backgrounds. **(C-E)** The bifurcation diagram for activated Thickveins (Tkv) **(C)**, nuclear pMad **(D)**, and Fu/Smurf **(E)** in *wild-type* and *dad^KO^* backgrounds.

We hypothesized that, under the model structure, it would be possible for the preCB to reside on the lower branch in wildtype, but on the higher branch in *dad* mutants. We used the steady states of the GSC-only model with *dad* removed as initial conditions. Under these conditions, we found that the CBs converged to the higher branch in 54% of the sets. Further analysis of these parameter sets showed that both cells begin with high levels of pMad, as the GSC-only steady state has a higher-than-wildtype concentration of pMad in *dad* mutants, as expected (Figure 7A-B). During cell division, as the mass-transfer coefficient decreases, pMad levels in the preCB decrease; however, the fractional decrease is less than that found in wildtype CBs. As such, pMad levels remain high for the entirety of the cell cycle in both the GSC and the preCB, and Fu/Smurf continues to be repressed throughout. Therefore, the steady state of active pMad from the preCB model often ends on the upper branch of the bifurcation diagram (Figure 7). Taken together, our analysis shows the dual NFL/PFL model structure allows Dad to control not only the levels of pMad in the GSC, but also the Fu/Smurf levels in, and hence, the differentiation of the CBs, even though Dad is not expressed, nor even ultimately present, in the CBs. As such, in wildtype simulations, Dad acts non-autonomously to ensure an initial condition for the CB that is consistent with a differentiation phenotype (Figure 8).

**Figure 8:**
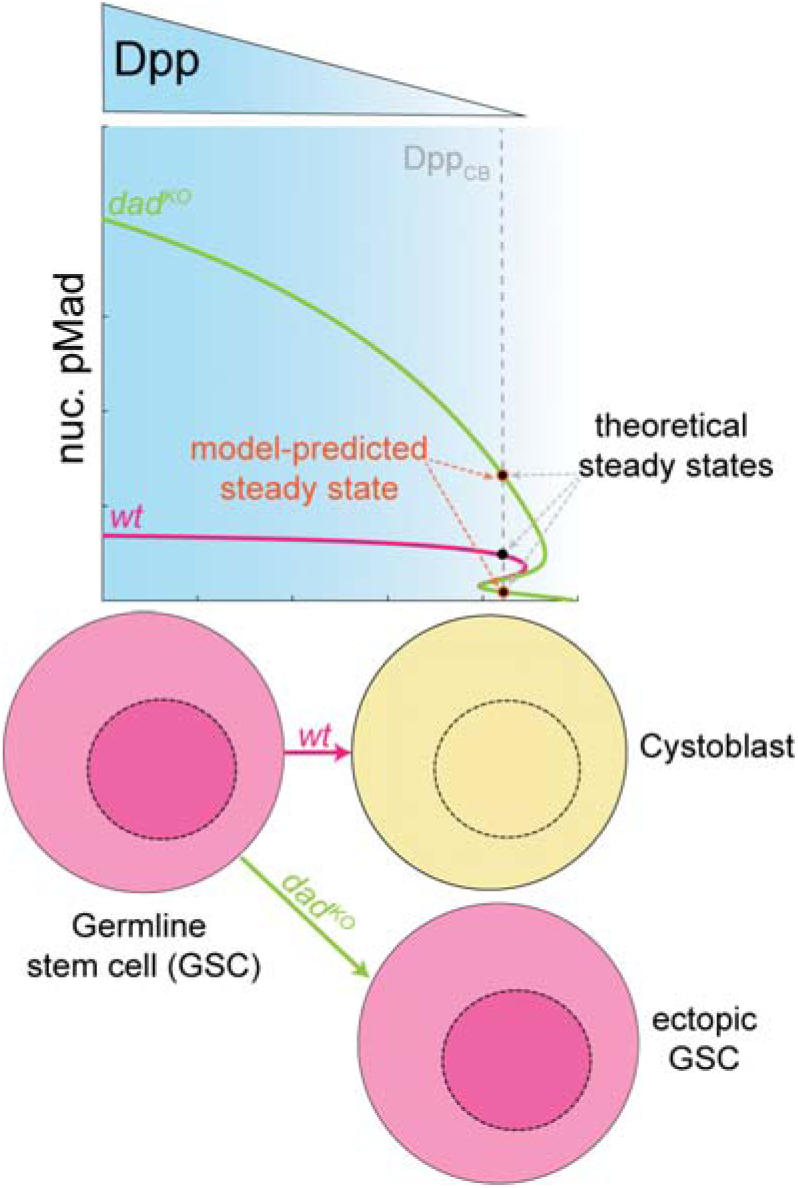
Dad acts non-autonomously to ensure Cystoblast differentiation. The negative feedback loop through Dad and the positive feedback loop through Fused allows for asymmetric divisions of

## 5 Discussion

The *Drosophila* germarium is a well-studied system to understand the dynamics of stem cell decisions (Drummond-Barbosa, 2019; Fisun Hamaratoglu and Markus Affolter and George, 2014; Fuller and Spradling, 2007; Harris and Ashe, 2011; Xie, 2013; Xie and Spradling, 2000; Yamashita, et al., 2005; Zhang and Kalderon, 2000). There are a several molecular tools available to endogenously manipulate the genome and quantify the dynamics of the transcripts and proteins *in vivo* in live tissues (Beumer, et al., 2013; Gratz, et al., 2015; Gratz, et al., 2014; Kanca, et al., 2019; Port, et al., 2014; Xue, et al., 2014; Yu, et al., 2013). However, a quantitative investigation of the factors which regulate stem cell homeostasis and differentiation is lacking.

Pargett et al. (2014) (Pargett, et al., 2014) analyzed qualitative data from the early studies of the germarium and proposed numerous mechanisms by developing predictive mathematical models. Since then, a few recent studies have examined the role of BMP signaling pathway in the germarium through transgenesis and advanced imaging techniques such as Fluorescence Recovery After Photobleaching (FRAP) to estimate protein mass transfer rates (Sardi, et al., 2021; Wilcockson and Ashe, 2021). This has facilitated a means to develop a mechanistic understanding of the role of the Dpp pathway in regulating stem cell behavior.

But overall, an understanding of the exact mechanism through which the BMP signal transduction pathway regulates stem cell decisions is lacking. The attempts have been limited due to requirement of long-time course imaging (Villa-Fombuena, et al., 2021), variability in the shape of the tissue, which impedes image analysis pipelines, and requirement of advanced imaging techniques such as high resolution microscopy to visualize stem cell activity over a 10 *μ*m region without photobleaching. Therefore, detailed mechanistic models are required to summarize data and to predict and design future experiments.

In this work, we developed a biologically-informed mathematical model to elucidate the role BMP pathway elements such as Dad (in negative feedback loop) and Fu/Smurf (in positive feedback loop) in regulating stem cell decisions. The model has four compartments to track the BMP signaling the two connected cells, GSC and the preCB, as proteins transport across the cells and into their nuclei. We evaluated the model predictions and constrained its performance by imposing constraints inferred from qualitative biological experiments.

Our analysis indicates that Dad renders robustness by modulating the concentration of transcriptionally active pMad/Medea in response to perturbations in the concentration of ligand-receptor complex and regulates rise time of Fu/Smurf in the CBs. In some parameter sets, the positive feedback loop through Fused enables the system to exhibit bistability, such that in a genetically *wildtype* background, the GSC starts with a high concentration of pMad/Medea, and over the course of division, the CB settles on the lower branch of pMad/Medea (upper branch of Fu/Smurf). Furthermore, the model shows a possible behavior in which, in a *dad*-removed background, the GSCs fail to modulate pMad/Medea robustly, allowing CBs to remain on the upper pMad branch and thereby failing to express high levels of Fu/Smurf. Physically, this translates to a hypothesis in which the initial levels of pMad in the pre-CB, inherited from the GSCs, determine whether the pre-CB ultimately differentiates. Moreover, these initial levels are determined by negative feedback through Dad expressed in the GSC.

However, our simulations predict that the frequency with which in a *wildtype* system the preCB lands on the lower branch (differentiation branch) is low, since it happens only in 10% of the simulations. It is known that several signaling pathways, including JAK/Stat and Hh, coordinate to enable CB de-differentiate to GSCs based on environmental cues to maintain the number of GSCs and CBs in the system (Liu, et al., 2015). Moreover, Pum-Brat mediated repression of Mad also renders bistability to the CBs (Harris, et al., 2011). Accounting for these factors in the mathematical model, would help explain why, in most parameter sets, the BMP pathway alone does not ensure that *wild-type* CBs differentiate.

The predictive power of this model can be further increased by performing live-cell imaging on the germarium *ex vivo* by generating endogenously tagged fluorescent reporter lines to Dpp pathway components such as Mad and Medea, and using the MS2-MCP (Bertrand, et al., 1998; Ferraro, et al., 2016; Forrest and Gavis, 2003; Garcia, et al., 2013; Lucas, et al., 2013) system to visualize nascent transcriptional bursts of *dad* in the germline to constrain the model mani-fold.

## 7 Data Availability Statement

The source code and data are uploaded here: https://github.com/gtreeves/GSC_feedback_loops.git

## 8 Acknowledgments

Portions of this research were conducted with the advanced computing resources provided by Texas A&M High Performance Research Computing.

This research was supported by NSF grant 2313692 awarded to G.T.R.

